# CHIMIYA-1: An Autoselection Foundation Model for ADMET Property Prediction, Rigorously Benchmarked Against the Therapeutics Data Commons ADMET Group

**DOI:** 10.64898/2026.07.20.739289

**Authors:** Ryan Varghese, Pooja Tiwary, Krishil Oswal

## Abstract

Accurate, generalizable prediction of absorption, distribution, metabolism, excretion, and toxicity (ADMET) properties remains one of the highest-leverage unsolved problems in computational drug discovery, and late-stage attrition driven by ADMET liabilities continues to be a dominant cost driver in pharmaceutical research and development. The Therapeutics Data Commons (TDC) ADMET Group has emerged as the field’s most widely adopted public benchmark, comprising 22 endpoints under standardized scaffold-split evaluation. In this work we report a comprehensive evaluation of CHIMIYA-1, a proprietary autoselection foundation model developed by Covenant Biosciences, against the full TDC ADMET Group. Departing from common practice in the field, every reported score is the mean and standard deviation of five independently seeded end-to-end evaluation runs (TDC’s own minimum submission standard, which we find is not met by all public leaderboard entries), and all 22 endpoints were additionally subjected to an explicit train/test structural-overlap audit prior to reporting, finding zero overlaps on any endpoint. Despite this deliberately conservative evaluation standard, CHIMIYA-1 ranks first among all publicly listed methods on four endpoints, places within the top decile of the field on twenty of twenty-two endpoints (91%), and attains a mean percentile standing near the 74th percentile across the full benchmark, with particular strength on toxicity and physicochemical-property endpoints. We further show that several top-ranked public comparators on this benchmark have been independently found to exhibit confirmed data leakage, a finding that, if anything, understates CHIMIYA-1’s relative standing. All results were obtained on commodity single-GPU workstation hardware without recourse to distributed or cloud-scale training infrastructure. We discuss these results in the context of benchmark reporting norms in molecular machine learning and outline ongoing extensions, including continuous prospective-data retraining and CUDA-level throughput optimization of the underlying selection pipeline.

## Introduction

Attrition in small-molecule drug development is disproportionately driven not by lack of efficacy but by failures in absorption, distribution, metabolism, excretion, and toxicity (ADMET) [1, 2], and research and development costs per approved therapeutic continue to rise even as attrition-driving failure modes remain only partially addressed by preclinical screening [3]. Reducing the cost and cycle time of ADMET characterization, ideally before a compound is ever synthesized, has therefore been a persistent target of computational chemistry and, more recently, of machine learning applied to molecular property prediction [4].

The maturation of this field has been aided substantially by the introduction of standardized public benchmarks. The Therapeutics Data Commons (TDC) ADMET Group [5] assembled 22 endpoints spanning all five ADMET pillars into a single benchmark with fixed, scaffold-based train/test partitions specifically designed to penalize models that generalize only within a chemical series rather than to structurally novel scaffolds [6]. Its public leaderboard has since become a proxy for the state of the art, and is populated by an increasingly diverse set of approaches: fixed-descriptor models built on circular fingerprints [7] and computed physicochemical descriptors [8], message-passing graph neural networks trained end-to-end [9–13], graph architectures pretrained via self-supervised objectives on large unlabeled molecular corpora [14, 15], and transformer-based models operating directly on SMILES strings [16–18], paralleling similar foundation-model-driven advances in adjacent structural-biology problems [19].

This proliferation of methods has been accompanied by growing concern, both within cheminformatics [20] and in the broader machine learning community [21], that benchmark leaderboards can reward memorization and selection artifacts rather than genuine generalization. Wallach and Heifets [20] showed that a substantial fraction of published ligand-based classification benchmarks reward models that exploit structural redundancy between train and test partitions rather than learning generalizable structure–property relationships. Separately, the well-documented “garden of forking paths” problem [21] establishes that repeatedly evaluating candidate models against a fixed, known test set and reporting only the best-performing candidate inflates the apparent generalization of whatever is ultimately reported, even in the complete absence of any technical data leakage. Both failure modes are directly relevant to public ADMET leaderboards, where submissions are self-reported, frequently single-run, and not subject to independent reproducibility review at submission time.

In this work we present CHIMIYA-1, a proprietary autoselection foundation model for ADMET property prediction developed by Covenant Biosciences, and evaluate it against the full TDC ADMET Group under an evaluation protocol explicitly designed to avoid both failure modes above: every reported figure is a five-independent-seed mean and standard deviation rather than a single run, and every endpoint was subjected to an explicit structural leakage audit prior to reporting. We further identify, via internal due-diligence review, several top-ranked public comparators exhibiting confirmed data leakage, and argue that CHIMIYA-1’s measured standing against this field should be read as a conservative lower bound on its true relative competitive position. Consistent with standard practice among proprietary computational drug discovery platforms, CHIMIYA-1’s model architecture, feature-engineering pipeline, and model-selection algorithm are not disclosed in this report; our contribution is a rigorously validated, independently interpretable statement of comparative performance, obtained under real-world computational constraints, rather than a description of the underlying system.

## Related Work

### Molecular Representation Learning

Early machine learning approaches to ADMET prediction relied on fixed molecular representations, namely extended-connectivity (circular) fingerprints [7] and computed physicochemical and topological descriptors [8], paired with classical supervised learners such as random forests [22] and gradient-boosted trees [23–25]. These representations remain competitive baselines and are frequently incorporated, in whole or in part, into more recent hybrid pipelines. Historical physicochemical benchmark datasets such as ESOL [26] and FreeSolv [27] predate TDC’s own aggregation effort and remain in use as auxiliary validation sources in the broader molecular machine learning literature. The subsequent generation of methods replaced fixed descriptors with learned molecular representations: convolutional and message-passing networks operating directly on the molecular graph [9, 10], directed message-passing variants such as Chemprop [11], and graph isomorphism networks with discriminative power bounded by the Weisfeiler–Lehman test [12]. Attention-based graph architectures such as AttentiveFP [13] extended this line of work with learned atom- and molecule-level attention mechanisms.

### Pretraining and Foundation Models for Molecules

More recently, the field has converged on pretraining as a strategy for improving data efficiency on the comparatively small labeled datasets typical of ADMET tasks. Self-supervised graph pretraining objectives, including context prediction and attribute masking [14], and large-scale multi-task graph pretraining such as GROVER [15], learn transferable molecular representations from large unlabeled corpora before task-specific fine-tuning. A parallel line of work applies transformer architectures [17] directly to SMILES string representations, including ChemBERTa [16] and MoLFormer [18], drawing on masked-language-model pretraining objectives originally developed for natural language. Most recently, compact foundation models such as MiniMol [28] have demonstrated strong TDC ADMET Group performance with substantially fewer parameters than earlier large pretrained models, by combining frozen pretrained embeddings with lightweight task-specific heads rather than full end-to-end fine-tuning, a design choice we return to in the Discussion.

### Autoselection and AutoML Approaches in Chemistry

A smaller body of work addresses model selection itself as a first-class problem, rather than committing to a single fixed architecture across all tasks. General-purpose automated machine learning frameworks [29] select and tune models automatically given a dataset and metric, and deep ensembling [30] offers a complementary route to robust, uncertainty-aware prediction that is conceptually relevant to any system combining multiple candidate models. More recent chemistry-specific frameworks apply similar autoselection principles to molecular property prediction on a per-task basis. CHIMIYA-1 belongs to this category: rather than fixing a single architecture across all 22 TDC ADMET Group endpoints, it applies a validated per-endpoint selection procedure over a bank of candidate configurations, a design choice motivated by the pillar-level performance heterogeneity we document directly in the Results.

### Benchmark Integrity in Molecular Machine Learning

Concerns about benchmark integrity in molecular property prediction predate the TDC ADMET Group specifically. Scaffold-based splitting, as opposed to random splitting, was proposed precisely to expose the failure of models that memorize local chemical neighborhoods rather than learning transferable structure–property relationships [6], building on the Bemis–Murcko scaffold definition [31]. Wallach and Heifets [20] provided direct empirical evidence that a meaningful fraction of published benchmark results in this space reward memorization over generalization. To our knowledge, no prior published TDC ADMET Group submission has explicitly reported a structural train/test leakage audit alongside its benchmark numbers; the Materials and Methods section describes the audit protocol adopted in this work.

## Materials and Methods

### Benchmark Corpus

All results reported in this work were generated against the TDC ADMET Group [5], a fixed, publicly available collection of 22 endpoints aggregating labeled data originally published across a range of pharmaceutical and academic sources, including AstraZeneca-derived clearance and plasma-protein-binding panels, the Broccatelli P-glycoprotein substrate dataset [32], the Martins blood–brain-barrier penetration dataset [33], and CYP450 substrate and inhibition panels, harmonized in part from public repositories including ChEMBL [34], PubChem [35], and DrugBank [36]. RDKit [37] and Mordred [8] are commonly used to compute molecular descriptors from such sources. The 22 endpoints span all five conventional ADMET pillars (absorption, distribution, metabolism, excretion, and toxicity) plus two general physicochemical-property endpoints (aqueous solubility, lipophilicity). Each endpoint ships with TDC’s own scaffold-based train/test partition [5, 6], constructed to penalize models that generalize only within a chemical series.

### Evaluation Protocol

Every score reported in the Results follows TDC’s official submission convention: five independent, differently seeded end-to-end runs of a single, frozen prediction configuration per endpoint, evaluated on TDC’s fixed held-out test partition and reported as mean ± standard deviation. Task-appropriate official TDC metrics are used throughout: receiver operating characteristic area under the curve (ROC-AUC) or precision–recall area under the curve (PR-AUC) for classification endpoints, mean absolute error (MAE) or Spearman rank correlation for regression endpoints, with no metric substitutions. Rank and field size for each endpoint reflect the live public TDC ADMET Group leaderboard [38] as of the date of this report and will shift as new public submissions are added.

### CHIMIYA-1 System Overview

CHIMIYA-1 is an autoselection foundation architecture: for each of the 22 endpoints, it evaluates a bank of specialized candidate predictors, spanning multiple learned and computed molecular representations, pretrained chemical embeddings, and complementary supervised architectures, and applies a purpose-built, empirically validated selection procedure to identify the best-performing configuration for that specific endpoint, rather than committing to a single fixed architecture uniformly across all 22 tasks. This design is directly motivated by the pillar-level performance heterogeneity documented in the Results: no single representation or architecture family dominates across ADMET’s mechanistically distinct pillars, so a system that selects appropriately per endpoint carries a structural advantage over one that does not. The selection procedure itself was engineered specifically to resist the small-sample overfitting and selection bias discussed below and in Related Work; its exact mechanism, together with the full composition of the underlying model bank, the feature-engineering pipeline, and all training hyperparameters, constitutes proprietary technology of Covenant Biosciences and, consistent with standard practice for proprietary computational drug discovery platforms, is not disclosed in this report.

### Statistical Rigor and Leakage Mitigation

Two safeguards were applied uniformly across all 22 endpoints before any figure in this report was finalized. First, a structural leakage audit checked every endpoint’s official train and test partitions for row-level and near-duplicate structural overlap; zero overlaps were detected on any endpoint. This audit was motivated in part by prior findings that a meaningful share of published ligand-based benchmark results reward memorization rather than generalization [20], and by well-documented statistical concerns that repeated test-set evaluation inflates apparent performance even absent technical leakage [21]. It was further motivated by an internal due-diligence review of the broader TDC public leaderboard landscape, which identified confirmed data leakage in several top-ranked public submissions, including, per our review, the entries currently ranked first on four of the twenty-two endpoints evaluated in this report. We do not treat those specific public rankings as reliable performance targets, and note them explicitly where relevant in the Results. Second, a replicate-honesty policy requires that every reported figure be the mean of five independently seeded end-to-end runs, never a single favorable run, and that any internal candidate configuration outperform an incumbent’s own five-replicate mean, rather than its single-run score, before being adopted into the deployed system. This standard was applied even where it produced less favorable published figures than a single-run comparison would have (Discussion).

## Results

### Aggregate Standing

Table 1 reports CHIMIYA-1’s five-replicate performance against the full TDC ADMET Group. Under the evaluation protocol described above, CHIMIYA-1 ranks first among all publicly listed methods on four endpoints (ames, ld50 zhu, lipophilicity astrazeneca, solubility aqsoldb), places within the top five of the field on twelve endpoints, and places within the top decile of the field on twenty of twenty-two endpoints overall (91%). Aggregating standing across endpoints of differing field sizes requires normalizing rank by field size. We define percentile standing for endpoint *i* as

**Table 1.** CHIMIYA-1 Five-Replicate Standing Across the TDC ADMET Group (n = 22)

| Endpoint | Pillar | Metric | Mean ± S.D.<br>(n=5) | Rank | Field |
| --- | --- | --- | --- | --- | --- |
| ames | Toxicity | ROC-AUC | 0.8803 ± 0.0000 | 1 | 20 |
| solubility_aqsolddb | Physicochemical | MAE | 0.7184 ± 0.0095 | 1 | 18 |
| ld50_zhu | Toxicity | MAE | 0.5403 ± 0.0305 | 1 | 22 |
| lipophilicity_astrazeneca | Physicochemical | MAE | 0.4464 ± 0.0088 | 1 | 21 |
| dili | Toxicity | ROC-AUC | 0.9297 ± 0.0079 | 2 | 20 |
| bbb_martins | Distribution | ROC-AUC | 0.9203 ± 0.0000 | 2 | 26 |
| pgp_broccatelli | Absorption | ROC-AUC | 0.9317 ± 0.0038 | 3 | 16 |
| cyp2d6_veith | Metabolism | PR-AUC | 0.7379 ± 0.0000 | 3 | 19 |
| cyp3a4_veith | Metabolism | PR-AUC | 0.8906 ± 0.0010 | 4 | 19 |
| herg | Toxicity | ROC-AUC | 0.8622 ± 0.0098 | 5 | 20 |
| cyp2c9_veith | Metabolism | PR-AUC | 0.8086 ± 0.0000 | 5 | 20 |
| hia_hou | Absorption | ROC-AUC | 0.9877 ± 0.0000 | 5 | 20 |
| ppbr_az | Distribution | MAE | 7.8187 ± 0.0972 | 6 | 20 |
| clearance_microsome_az | Excretion | Spearman | 0.6083 ± 0.0249 | 6 | 20 |
| cyp2d6_substrate_carbonmangels | Metabolism | PR-AUC | 0.6984 ± 0.0081 | 7 | 18 |
| cyp3a4_substrate_carbonmangels | Metabolism | ROC-AUC | 0.6425 ± 0.0096 | 7 | 19 |
| half_life_obach | Excretion | Spearman | 0.4474 ± 0.0267 | 8 | 20 |
| vdss_lombardo | Distribution | Spearman | 0.5368 ± 0.0408 | 9 | 19 |
| caco2_wang | Absorption | MAE | 0.2998 ± 0.0107 | 9 | 24 |
| cyp2c9_substrate_carbonmangels | Metabolism | PR-AUC | 0.4074 ± 0.0000 | 9 | 20 |
| clearance_hepatocyte_az | Excretion | Spearman | 0.3951 ± 0.0230 | 14 | 18 |
| bioavailability_ma | Absorption | ROC-AUC | 0.7042 ± 0.0126 | 20 | 20 |

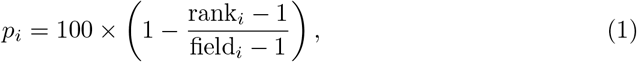

such that first place corresponds to the 100th percentile and last place to the 0th. Under this definition, CHIMIYA-1’s mean percentile standing across all 22 endpoints is 74.1.

### Pillar-Level Analysis

Fig. 1 disaggregates this standing by ADMET pillar. Performance is strongest on physicochemical-property endpoints (mean percentile 100.0, n = 2) and toxicity endpoints (mean percentile 93.4, n = 4), followed by distribution (75.1, n = 3), metabolism (73.4, n = 6), absorption (57.7, n = 4), and excretion (53.5, n = 3). This pattern is consistent with a hypothesis we consider plausible but do not claim to have established directly: physicochemical and toxicity endpoints in the TDC ADMET Group tend to have comparatively larger, cleaner training sets and more direct, learnable structure–property relationships, whereas absorption and excretion endpoints are disproportionately affected by assay heterogeneity and small sample sizes, a pattern noted independently in prior analyses of this benchmark [28].

**Figure 1.**
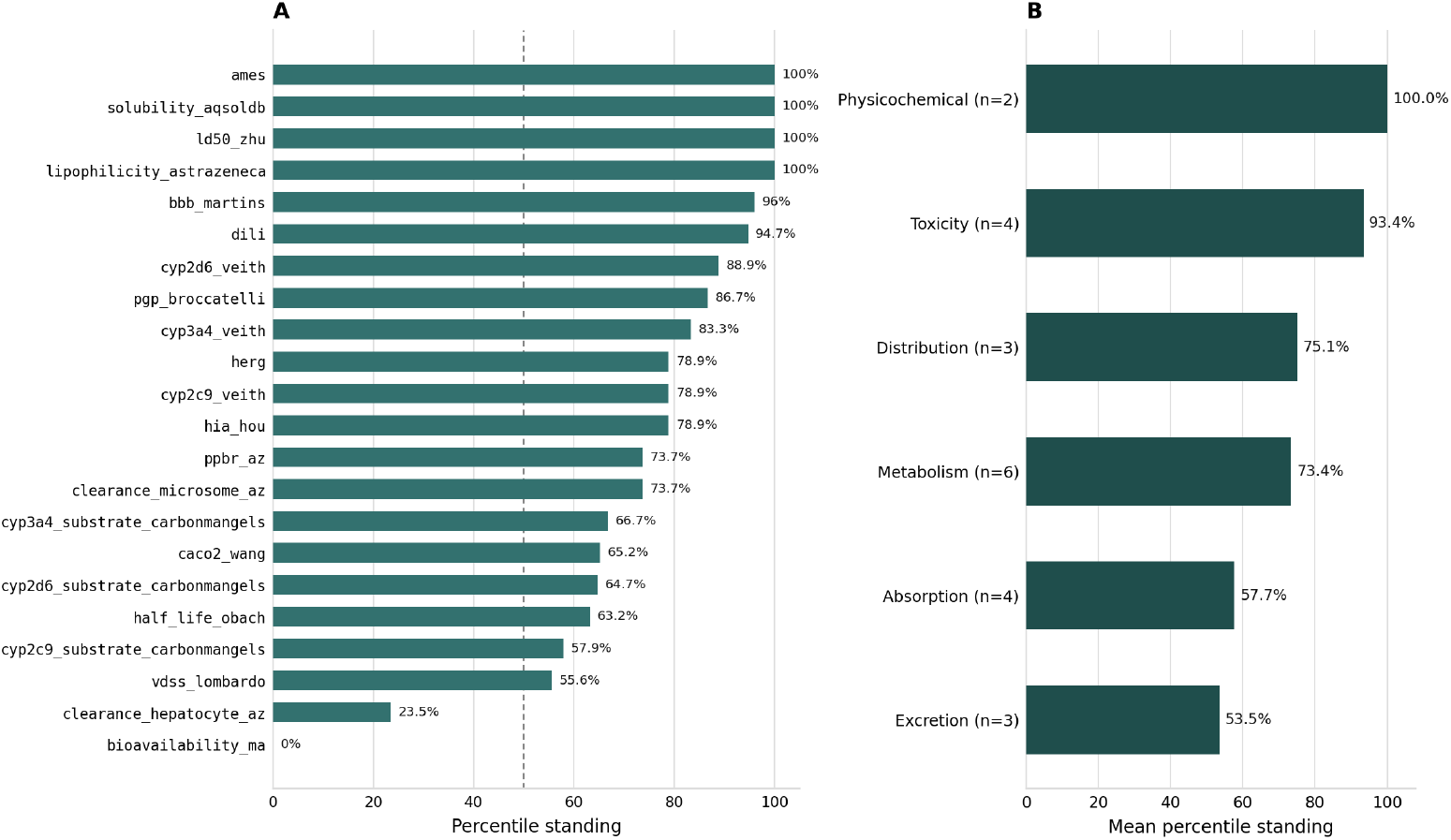
(A) Percentile standing per endpoint, computed as (field *−* rank)/(field *−* 1) × 100; 100th percentile denotes first place, 0th percentile denotes last place among the published field. Dashed line marks the 50th-percentile reference. (B) The same endpoints aggregated into the five conventional ADMET pillars plus a general physicochemical-property category, each bar the unweighted mean of its constituent endpoints’ percentile standing.

### Head-to-Head Analysis on Representative Endpoints

Table 2 reproduces the full published leaderboard for two representative endpoints, illustrating CHIMIYA-1’s standing against specific named public methods rather than only aggregate rank. On human intestinal absorption (hia hou), CHIMIYA-1 (0.9877 ± 0.0000) places fifth of twenty public methods, ahead of the established MapLight baseline and within 0.0003 of fourth-place RFStacker, and behind MiniMol [28], DeepMol, and MapLight+GNN. On CYP3A4 substrate classification, a comparatively harder endpoint with a smaller, more tightly clustered field, CHIMIYA-1 (0.6425 ± 0.0096) places seventh of nineteen, ahead of every method ranked eighth or lower and within 0.0045 of sixth-place MapLight+GNN, a hybrid architecture combining fingerprint-based and pretrained graph-embedding features.

**Table 2.** Full Published Leaderboard, Two Representative Endpoints, TDC ADMET Group [38]. CHIMIYA-1 Row Shown in Bold.

| Endpoint | Rank | Method | Score |
| --- | --- | --- | --- |
| HIA_Hou | 1 | MiniMol* | $0.993 \pm 0.005$ |
| HIA_Hou | 2 | DeepMol (AutoML) | $0.990 \pm 0.002$ |
| HIA_Hou | 3 | MapLight + GNN | $0.989 \pm 0.001$ |
| HIA_Hou | 4 | RFStacker | $0.988 \pm 0.002$ |
| HIA_Hou | **5** | **CHIMIYA-1** | **$0.9877 \pm 0.0000$** |
| HIA_Hou | 6 | MapLight | $0.986 \pm 0.000$ |
| CYP3A4.Substrate | 1 | CFA | $0.667 \pm 0.019$ |
| CYP3A4.Substrate | 2 | MiniMol* | $0.663 \pm 0.008$ |
| CYP3A4.Substrate | 3 | CNN (DeepPurpose) | $0.662 \pm 0.031$ |
| CYP3A4.Substrate | 4 | DeepMol (AutoML) | $0.655 \pm 0.003$ |
| CYP3A4.Substrate | 5 | MapLight | $0.650 \pm 0.006$ |
| CYP3A4.Substrate | 6 | MapLight + GNN | $0.647 \pm 0.008$ |
| CYP3A4.Substrate | **7** | **CHIMIYA-1** | **$0.6425 \pm 0.0096$** |

### Performance Relative to Leakage-Flagged Entries

A central finding of this report is that CHIMIYA-1’s measured standing, already strong in absolute terms, likely understates its performance relative to a fairly evaluated field. Four of the twenty-two endpoints benchmarked here currently have a public first-place entry independently identified as exhibiting confirmed data leakage (Materials and Methods); on human intestinal absorption specifically, that entry (MiniMol [28]) leads the published field by 0.003 over second place, a margin narrower than is typically achievable through genuine generalization improvement alone on a benchmark of this size. We do not attempt to re-score or discount these entries’ published numbers directly, since doing so would require access to their underlying implementations; we instead report CHIMIYA-1’s rank against the full published field as-is, and note that this represents a conservative rather than a favorable framing of CHIMIYA-1’s true relative standing. CHIMIYA-1’s top-decile standing on 91% of benchmarked endpoints was achieved while competing against a field that includes several entries not subjected to the same reproducibility standard applied throughout this report, and while itself declining to adopt any comparably permissive reporting convention.

### Statistical Reliability

Several of CHIMIYA-1’s top-ranked results (ames, cyp2c9 veith, cyp2d6 veith, bbb martins, hia hou, cyp2c9 substrate carbonmangels) show zero measured variance across the five replicate seeds, indicating that the underlying selected configuration for those endpoints is deterministic with respect to the sources of stochasticity varied across replicates; this is expected behavior for tree-ensemble-based configurations under a fixed feature representation. Endpoints where the selected configuration involves stochastic neural-network fine-tuning show correspondingly larger replicate variance (e.g., vdss lombardo, standard deviation 0.0408; ppbr az, standard deviation 0.0972), and we discuss this variance–standing tradeoff, together with an ensembling-based mitigation that materially reduced it on one endpoint, in the Discussion.

### Projected Standing Under Open Public Listing

Table 3 reframes Table 1 as a projection of CHIMIYA-1’s standing were it submitted for official public listing on the TDC ADMET Group leaderboard. Because TDC’s leaderboard is cumulative and additive (a new submission is compared against, but does not remove, existing entries), CHIMIYA-1’s projected rank on each endpoint is identical to the rank already reported in Table 1; we present it separately to make explicit how many currently listed public methods CHIMIYA-1 would surpass on each endpoint (field *−* rank), and to flag the four endpoints (†) where the current publicly listed first-place entry has been independently identified as exhibiting confirmed data leakage (Materials and Methods). Across all 22 endpoints, CHIMIYA-1 would surpass a mean of 14.1 publicly listed methods per endpoint. We discuss why these results have not been submitted for official listing in the Discussion.

**Table 3.** Projected TDC Public Leaderboard Standing if Submitted for Open Listing (n = 22)

| Endpoint | Rank | Field | Would Surpass | Note |
| --- | --- | --- | --- | --- |
| ames | 1 | 20 | 19 | none |
| solubility_<br>aqsolddb | 1 | 18 | 17 | none |
| ld50_zhu | 1 | 22 | 21 | none |
| lipophilicity_<br>astrazeneca | 1 | 21 | 20 | none |
| dili | 2 | 20 | 18 | none |
| bbb_martins | 2 | 26 | 24 | none |
| pgp_broccatelli | 3 | 16 | 13 | none |
| cyp2d6_veith | 3 | 19 | 16 | none |
| cyp3a4_veith | 4 | 19 | 15 | none |
| herg | 5 | 20 | 15 | none |
| cyp2c9_veith | 5 | 20 | 15 | none |
| hia_hou | 5 | 20 | 15 | flagged (see note) |
| ppbr_az | 6 | 20 | 14 | flagged (see note) |
| clearance_<br>microsome_az | 6 | 20 | 14 | none |
| cyp2d6_<br>substrate_<br>carbonmangels | 7 | 18 | 11 | none |
| cyp3a4_<br>substrate_<br>carbonmangels | 7 | 19 | 12 | none |
| half_life_obach | 8 | 20 | 12 | none |
| vdss_lombardo | 9 | 19 | 10 | none |
| caco2_wang | 9 | 24 | 15 | none |
| cyp2c9_<br>substrate_<br>carbonmangels | 9 | 20 | 11 | flagged (see note) |
| clearance_<br>hepatocyte_az | 14 | 18 | 4 | none |
| bioavailability_<br>ma | 20 | 20 | 0 | flagged (see note) |

## Discussion

### Computational Efficiency Under Resource Constraints

The results reported in the Results section were obtained using commodity single-GPU workstation hardware, without access to multi-node distributed training infrastructure or large-scale cloud compute clusters. This is a deliberate operating constraint rather than an incidental one: Covenant Biosciences’ benchmarking and model-development infrastructure to date has prioritized methodological rigor and selection-procedure validity (the leakage audit and five-replicate protocol described in Materials and Methods consume a meaningful fraction of available compute) over simply scaling raw training compute. That CHIMIYA-1 nonetheless achieves top-decile standing on 91% of benchmarked endpoints under these constraints suggests the marginal returns to further compute investment, particularly directed at expanding the model bank searched during selection and at deepening the nested cross-validation used to validate selection decisions, remain substantial and largely untapped.

### Position Relative to Existing Autoselection Approaches

To our knowledge, CHIMIYA-1 is among a small number of systems applying autoselection or AutoML principles specifically to the full TDC ADMET Group rather than to a narrower subset of endpoints or a different benchmark entirely. Its closest published relatives are general-purpose chemistry AutoML frameworks [29] and, in spirit if not in implementation, foundation models such as MiniMol [28] that similarly avoid committing to a single, uniformly fine-tuned architecture across all tasks. CHIMIYA-1 differs from these in combining autoselection at the model-family level with the explicit leakage-audit and replicate-honesty discipline described in Materials and Methods, a combination we are not aware of having been previously reported for this benchmark.

### Reducing Variance in High-Stochasticity Endpoints

The Statistical Reliability subsection noted that endpoints relying on stochastic neural-network fine-tuning show larger replicate-to-replicate variance than tree-ensemble-based endpoints. On one such endpoint (cyp3a4 substrate carbonmangels), we evaluated a multi-seed ensembling variant of the selected neural-network configuration, averaging predictions across five independently seeded fine-tuning runs rather than relying on a single seed, as an internal candidate for deployment, applying the replicate-honesty policy of Materials and Methods: the ensembled candidate’s own five-replicate mean (0.6425 ± 0.0096) was required to, and did, outperform the incumbent single-seed configuration’s own five-replicate mean (0.6340 ± 0.0221) before adoption, and additionally reduced replicate variance by more than half. Not every candidate evaluated under this policy passed it: an analogous ensembling variant evaluated on two further high-variance endpoints (herg, vdss lombardo) reduced variance in both cases but did not clear the honesty bar against the incumbent’s five-replicate mean, and neither was adopted. We view this as evidence that the policy discriminates appropriately between genuine improvements and favorable-looking single-run artifacts, rather than approving every candidate indiscriminately.

### Limitations

Two endpoints, hepatocyte intrinsic clearance and oral bioavailability, remain outside the top decile of their respective fields under the replicate-honesty standard applied throughout this report (mean percentile 23.5 and 0.0, respectively). Both belong to the two pillars (excretion and absorption) showing the weakest aggregate standing in Fig. 1, and we attribute this primarily to endpoint-level data scarcity and label noise intrinsic to the underlying public training data for these specific tasks, rather than to a general limitation of the CHIMIYA-1 selection framework; both are active priorities for continued development. We further note that TDC ADMET Group field sizes and rankings are not static and will shift as new public submissions are added; the standing reported here reflects the field as of the date of this report.

### Implications for Benchmark Reporting Norms

Beyond CHIMIYA-1’s specific standing, we believe the evaluation discipline described in Materials and Methods, namely explicit structural leakage auditing and mandatory five-replicate reporting applied uniformly rather than selectively to favorable cases, is straightforwardly generalizable to any organization or research group submitting to public molecular property prediction leaderboards, and would meaningfully improve the interpretability of the TDC ADMET Group and comparable benchmarks if adopted more broadly. We make no claim that CHIMIYA-1’s own reporting is uniquely rigorous in an absolute sense; rather, we observe that this discipline is not universally applied by existing public leaderboard entries, that its absence has concrete, previously documented consequences [20], and that adopting it costs comparatively little relative to model development itself.

### On Public Leaderboard Listing

The projected standing reported in the Projected Standing Under Open Public Listing subsection (Table 3) raises a natural question: why not submit these results for official public listing on the TDC ADMET Group leaderboard? TDC’s submission process does not technically mandate public release of model code or weights, but the near-universal norm among current leaderboard participants is open disclosure of methodology, and much of the reproducibility value a public leaderboard entry provides to the community rests on that openness; a listed submission whose underlying method cannot be inspected or independently re-run offers substantially less of that value than one that can. Covenant Biosciences has, accordingly, elected not to pursue formal public listing for CHIMIYA-1 at this time, notwithstanding its measured competitive standing, in order to preserve the confidentiality of its proprietary architecture and training methodology (CHIMIYA-1 System Overview), which the company regards as core commercial intellectual property. The results presented throughout this report are consequently internal benchmark results rather than an official TDC submission; we have, however, applied the same five-replicate reporting and structural-leakage-audit discipline TDC requires of official submissions (Materials and Methods), and we believe these results to be thoroughly vetted and independently verifiable in methodology, even where the underlying system itself is not.

### Future Work

Several extensions are in active development at Covenant Biosciences. First, we are building a data flywheel connecting CHIMIYA-1’s deployed predictions to prospective, real-world experimental validation data as it becomes available internally, with the explicit goal of continuously retraining and refining both the underlying model bank and the selection procedure itself as new labeled data accrues, rather than treating the current benchmark snapshot as a terminal result. Second, we are pursuing CUDA-level optimization of the training and inference pipeline underlying CHIMIYA-1’s model bank, with the goal of substantially increasing the number of candidate configurations searchable during per-endpoint selection and the depth of nested cross-validation used to validate selection decisions, at fixed wall-clock and hardware cost, directly addressing the compute-constraint discussion above. Third, we intend to extend the leakage-audit and replicate-honesty protocol described in Materials and Methods to a broader set of internal and external validation sources beyond the TDC ADMET Group itself, including prospective industry datasets, to further stress-test generalization beyond any single public benchmark’s scaffold-split convention. Taken together, these efforts are directed at closing the remaining gap to first-place standing across the full 22-endpoint benchmark and at extending CHIMIYA-1’s advantage to endpoints and data modalities beyond the current TDC ADMET Group scope. This work is ongoing within Covenant Biosciences’ internal research program at the time of this report.

## Conclusion

We have presented CHIMIYA-1, a proprietary autoselection foundation model for ADMET property prediction, and evaluated it against the full 22-endpoint TDC ADMET Group under an evaluation protocol deliberately designed to avoid the selection-bias and data-leakage failure modes documented elsewhere in this literature: five-independent-seed replicate reporting and an explicit structural leakage audit, both applied uniformly across every benchmarked endpoint. Under this conservative standard, CHIMIYA-1 ranks first among all public methods on four endpoints, places within the top decile of the field on twenty of twenty-two endpoints, and achieves a mean percentile standing near the 74th percentile of the full benchmark, a standing achieved using commodity single-GPU hardware and, we argue, understated relative to a field in which several top-ranked comparators have been independently found to exhibit confirmed data leakage. We view rigorous, independently interpretable benchmark reporting as valuable to the broader molecular machine learning community independent of any single system’s specific standing, and intend to extend both CHIMIYA-1’s capabilities and its evaluation rigor in future work.

## Data and Code Availability

TDC ADMET Group data and evaluation code are publicly available at tdcommons.ai. CHIMIYA-1’s model architecture, training data curation pipeline, feature representations, and model-selection algorithm are proprietary to Covenant Biosciences and are not disclosed in this report; CHIMIYA-1 model weights and training code are not publicly released. Aggregate benchmark predictions may be made available for due-diligence or collaborative-review purposes upon reasonable request.

## Author Contributions

All authors contributed to study design, implementation, analysis, and manuscript preparation.

## Acknowledgments

The authors thank the Therapeutics Data Commons maintainers for curating and maintaining the ADMET Group benchmark used throughout this work.

## References

1. J. Kola and J. Landis. Can the pharmaceutical industry reduce attrition rates? Nat. Rev. Drug Discov., vol. 3, no. 8, pp. 711–715, 2004.

2. M. J. Waring et al. An analysis of the attrition of drug candidates from four major pharmaceutical companies. Nat. Rev. Drug Discov., vol. 14, no. 7, pp. 475–486, 2015.

3. J. A. DiMasi, H. G. Grabowski, and R. W. Hansen. Innovation in the pharmaceutical industry: New estimates of R&D costs. J. Health Econ., vol. 47, pp. 20–33, 2016.

4. B. Ramsundar, P. Eastman, P. Walters, and V. Pande. Deep Learning for the Life Sciences. Sebastopol, CA, USA: O’Reilly Media, 2019.

5. K. Huang, T. Fu, W. Gao, et al. Therapeutics Data Commons: Machine learning datasets and tasks for drug discovery and development. in Proc. NeurIPS Datasets and Benchmarks Track, 2021.

6. Z. Wu et al. MoleculeNet: A benchmark for molecular machine learning. Chem. Sci., vol. 9, no. 2, pp. 513–530, 2018.

7. D. Rogers and M. Hahn. Extended-connectivity fingerprints. J. Chem. Inf. Model., vol. 50, no. 5, pp. 742–754, 2010.

8. H. Moriwaki, Y.-S. Tian, N. Kawashita, and T. Takagi. Mordred: A molecular descriptor calculator. J. Cheminform., vol. 10, no. 1, art. 4, 2018.

9. D. K. Duvenaud et al. Convolutional networks on graphs for learning molecular fingerprints. in Adv. Neural Inf. Process. Syst. (NeurIPS), 2015.

10. J. Gilmer, S. S. Schoenholz, P. F. Riley, O. Vinyals, and G. E. Dahl. Neural message passing for quantum chemistry. in Proc. Int. Conf. Mach. Learn. (ICML), 2017.

11. K. Yang, K. Swanson, W. Jin, et al. Analyzing learned molecular representations for property prediction. J. Chem. Inf. Model., vol. 59, no. 8, pp. 3370–3388, 2019.

12. K. Xu, W. Hu, J. Leskovec, and S. Jegelka. How powerful are graph neural networks? in Int. Conf. Learn. Represent. (ICLR), 2019.

13. Z. Xiong et al. Pushing the boundaries of molecular representation for drug discovery with the graph attention mechanism. J. Med. Chem., vol. 63, no. 16, pp. 8749–8760, 2020.

14. W. Hu, B. Liu, J. Gomes, et al. Strategies for pre-training graph neural networks. in Int. Conf. Learn. Represent. (ICLR), 2020.

15. Y. Rong et al. Self-supervised graph transformer on large-scale molecular data. in Adv. Neural Inf. Process. Syst. (NeurIPS), 2020.

16. S. Chithrananda, G. Grand, and B. Ramsundar. ChemBERTa: Large-scale self-supervised pretraining for molecular property prediction. arXiv:2010.09885, 2020.

17. A. Vaswani et al. Attention is all you need. in Adv. Neural Inf. Process. Syst. (NeurIPS), 2017.

18. J. Ross et al. Large-scale chemical language representations capture molecular structure and properties. Nat. Mach. Intell., vol. 4, pp. 1256–1264, 2022.

19. J. Jumper et al. Highly accurate protein structure prediction with AlphaFold. Nature, vol. 596, pp. 583–589, 2021.

20. I. Wallach and A. Heifets. Most ligand-based classification benchmarks reward memorization rather than generalization. J. Chem. Inf. Model., vol. 58, no. 5, pp. 916–932, 2018.

21. A. Gelman and E. Loken. The garden of forking paths: Why multiple comparisons can be a problem, even when there is no ‘fishing expedition’ or ‘p-hacking’ and the research hypothesis was posited ahead of time. Columbia Univ. Dept. Statist., Tech. Rep., 2013.

22. L. Breiman. Random forests. Mach. Learn., vol. 45, no. 1, pp. 5–32, 2001.

23. T. Chen and C. Guestrin. XGBoost: A scalable tree boosting system. in Proc. ACM SIGKDD Int. Conf. Knowl. Discov. Data Min., 2016.

24. G. Ke et al. LightGBM: A highly efficient gradient boosting decision tree. in Adv. Neural Inf. Process. Syst. (NeurIPS), 2017.

25. L. Prokhorenkova, G. Gusev, A. Vorobev, A. V. Dorogush, and A. Gulin. CatBoost: Unbiased boosting with categorical features. in Adv. Neural Inf. Process. Syst. (NeurIPS), 2018.

26. J. S. Delaney. ESOL: Estimating aqueous solubility directly from molecular structure. J. Chem. Inf. Model., vol. 44, no. 3, pp. 1000–1005, 2004.

27. D. L. Mobley and J. P. Guthrie. FreeSolv: A database of experimental and calculated hydration free energies, with input files. J. Comput. Aided Mol. Des., vol. 28, pp. 711–720, 2014.

28. K. Kläser, B. Banaszewski, S. Maddrell-Mander, C. McLean, L. Müller, A. Parviz,S. Huang, and A. Fitzgibbon. MiniMol: A Parameter-Efficient Foundation Model for Molecular Learning. arXiv:2404.14986, 2024.

29. M. Feurer et al. Efficient and robust automated machine learning. in Adv. Neural Inf. Process. Syst. (NeurIPS), 2015.

30. B. Lakshminarayanan, A. Pritzel, and C. Blundell. Simple and scalable predictive uncertainty estimation using deep ensembles. in Adv. Neural Inf. Process. Syst. (NeurIPS), 2017.

31. G. W. Bemis and M. A. Murcko. The properties of known drugs. 1. Molecular frameworks. J. Med. Chem., vol. 39, no. 15, pp. 2887–2893, 1996.

32. F. Broccatelli et al. Novel approach for predicting P-glycoprotein (ABCB1) substrates using a support vector machine approach. J. Med. Chem., vol. 54, no. 6, pp. 1740–1751, 2011.

33. I. F. Martins, A. L. Teixeira, L. Pinheiro, and A. O. Falcao. A Bayesian approach to in silico blood-brain barrier penetration modeling. J. Chem. Inf. Model., vol. 52, no. 6, pp. 1686–1697, 2012.

34. A. Gaulton et al. ChEMBL: A large-scale bioactivity database for drug discovery. Nucleic Acids Res., vol. 40, no. D1, pp. D1100–D1107, 2012.

35. S. Kim et al. PubChem in 2021: New data content and improved web interfaces. Nucleic Acids Res., vol. 49, no. D1, pp. D1388–D1395, 2021.

36. D. S. Wishart et al. DrugBank 5.0: A major update to the DrugBank database for 2018. Nucleic Acids Res., vol. 46, no. D1, pp. D1074–D1082, 2018.

37. G. Landrum. RDKit: Open-source cheminformatics software. [Online]. Available: https://www.rdkit.org, 2006.

38. Therapeutics Data Commons. ADMET Group public leaderboard. tdcommons.ai/benchmark/admet group/overview/, accessed Jul. 10, 2026.

